# Region-specific expression of young small-scale duplications in the human central nervous system

**DOI:** 10.1101/821256

**Authors:** Solène Brohard-Julien, Vincent Frouin, Vincent Meyer, Smahane Chalabi, Jean-François Deleuze, Edith Le Floch, Christophe Battail

## Abstract

**Background:** The duplication of genes is one of the main genetic mechanisms that led to the gain in complexity of biological tissue. Although the implication of duplicated gene expression in brain evolution was extensively studied through comparisons between organs, their role in the regional specialization of the adult human central nervous system has not yet been well described.

**Results:** Our work explored intra-organ expression properties of paralogs through multiple territories of the human central nervous system (CNS) using transcriptome data generated by the Genotype-Tissue Expression (GTEx) consortium. Interestingly, we found that paralogs were associated with region-specific expression in CNS, suggesting their involvement in the differentiation of these territories. Beside the influence of gene expression level on region-specificity, we observed the contribution of both duplication age and duplication type to the CNS region-specificity of paralogs. Indeed, we found that small scale duplicated genes (SSDs) and in particular ySSDs (SSDs younger than the 2 rounds of whole genome duplications) were more CNS region-specific than other paralogs. Next, by studying the two paralogs of ySSD pairs, we observed that when they were region-specific, they tend to be specific to the same region more often than for other paralogs, showing the high co-expression of ySSD pairs. Extension of this analysis to families of paralogs showed that the families with co-expressed gene members (i.e. homogeneous families) were enriched in ySSDs. Furthermore, these homogeneous families tended to be region-specific families, where the majority of their gene members were specifically expressed in the same region.

**Conclusions:** Overall, our study suggests the major involvement of ySSDs in the differentiation of human central nervous system territories. Therefore, we show the relevance of exploring region-specific expression of paralogs at the intra-organ level.

## BACKGROUND

Comparative genomics and large-scale transcriptional studies have highlighted the major contribution of gene duplication to tissue differentiation and phenotypic diversity (1,2). The fact that some paralogs are retained in genomes through evolution seems to be initially favored by dosage balance (3) and their long-term preservation is then made possible by the following two processes: the neo-functionalization, which consists of the gain of a new function by one duplicate potentially associated with a different spatial expression (4–7), or the sub-functionalization which consists in the partition of the ancestral function or spatial expression between duplicates (8,9). The divergence of spatial expression between paralogs can be studied by the analysis of broad or narrow gene expression patterns across a collection of tissues (3,10,11). The comparison of transcriptomes between different mouse organs has shown that the brain was among the ones expressing the highest proportion of tissue-specific paralogs in relation to the total number of genes expressed in the brain, while it does not express the highest proportion of tissue-specific singletons (10). The brain is therefore a model perfectly suited to explore the intra-organ expression heterogeneity of the duplicated genes.

Among the 60% of human genes considered as paralogs (2), some come from whole-genome duplications (WGD) in early vertebrate lineage approximately 500 million years ago (12,13), the others come from small scale duplications (SSD) that have occurred throughout the evolution (14). A comparison in mammals, notably in humans, of the brain transcriptome with those of other organs has shown that WGDs tend to be enriched in brain-specific genes compared to SSDs (15,16). This supports the theory that genome duplications have allowed vertebrates to develop more complex cellular organizations of the central nervous system (CNS) (17,18).

In complement of the role of the WGDs in the tissue complexity, some studies support the idea that young duplicated genes tend to be preferentially expressed in evolutionarily young tissues. A higher proportion of primate-specific paralogs were found to be up-regulated in the developing human brain compared to the adult brain (19), whereas this expression pattern was not found for older duplications (20). Regarding recent duplications, that emerged in the human lineage, studies have suggested their contribution to human-specific adaptive traits, such as the gain of brain complexity (21–23).

While the expression properties of paralogs between different organs, including the brain, have been well studied, we have little knowledge of the expression characteristics of duplicated genes between different regions of the same organ. Large-scale transcriptional profiling of neuroanatomic regions (24) allows us now to further investigate paralog expression between the different territories of the human CNS according to their evolutionary properties. Exploring gene expression in this frame of reference, restricted to the CNS territories, makes it possible to identify distinct gene features which could be masked by transcriptome comparisons performed across several organs.

This present study explores in detail the expression patterns of paralogs between the different territories of the human CNS, using the GTEx resource, according to their evolutionary characteristics and gene families. We started assessing the changes in expression of duplicated genes between CNS regions and investigating paralogs expressed specifically in certain regions. Secondly, we studied the evolutionary characteristics of these paralogs with regional expression such as their age and the type of duplication event. We then analyzed the organization of paralogs in families using co-expression and studied their CNS region-specificity and their evolutionary characteristics.

## RESULTS

### 1/ Association of paralog expression with CNS differentiation

We considered in our study all human protein coding genes and the information collected on duplication events in order to split the gene population into paralogs and singletons (2) (Methods). In a landmark contribution, the GTEx (Genotype-Tissue Expression) consortium used RNA sequencing technology to establish the landscape of human gene expression across a large collection of postmortem biopsies (24). Gene expression data for hundreds of individuals from 13 normal brain-related regions (Methods) were obtained from the GTEx consortium. After filtering out low information content genes (Methods), abundance values of 16,427 protein-coding genes, including 10,335 paralogs and 6,092 singletons were retained.

Previous work by GTEx established the relevance of using gene expression data to cluster samples obtained from the same organ or from the same region of the CNS, even though assigning samples to the correct CNS region was more difficult (24). We focused specifically on CNS regions and assessed whether paralog expression could better classify samples into regions than the other genes. The unsupervised hierarchical classification of human CNS samples, based on their pair-wise similarity in terms of correlation across paralog expression values, was able to group together most samples belonging to the same region (Methods; Fig. 1). We observed a similar CNS region classification considering all protein-coding genes or only singletons (Additional File 1:Figure. S1).

**Figure 1.**
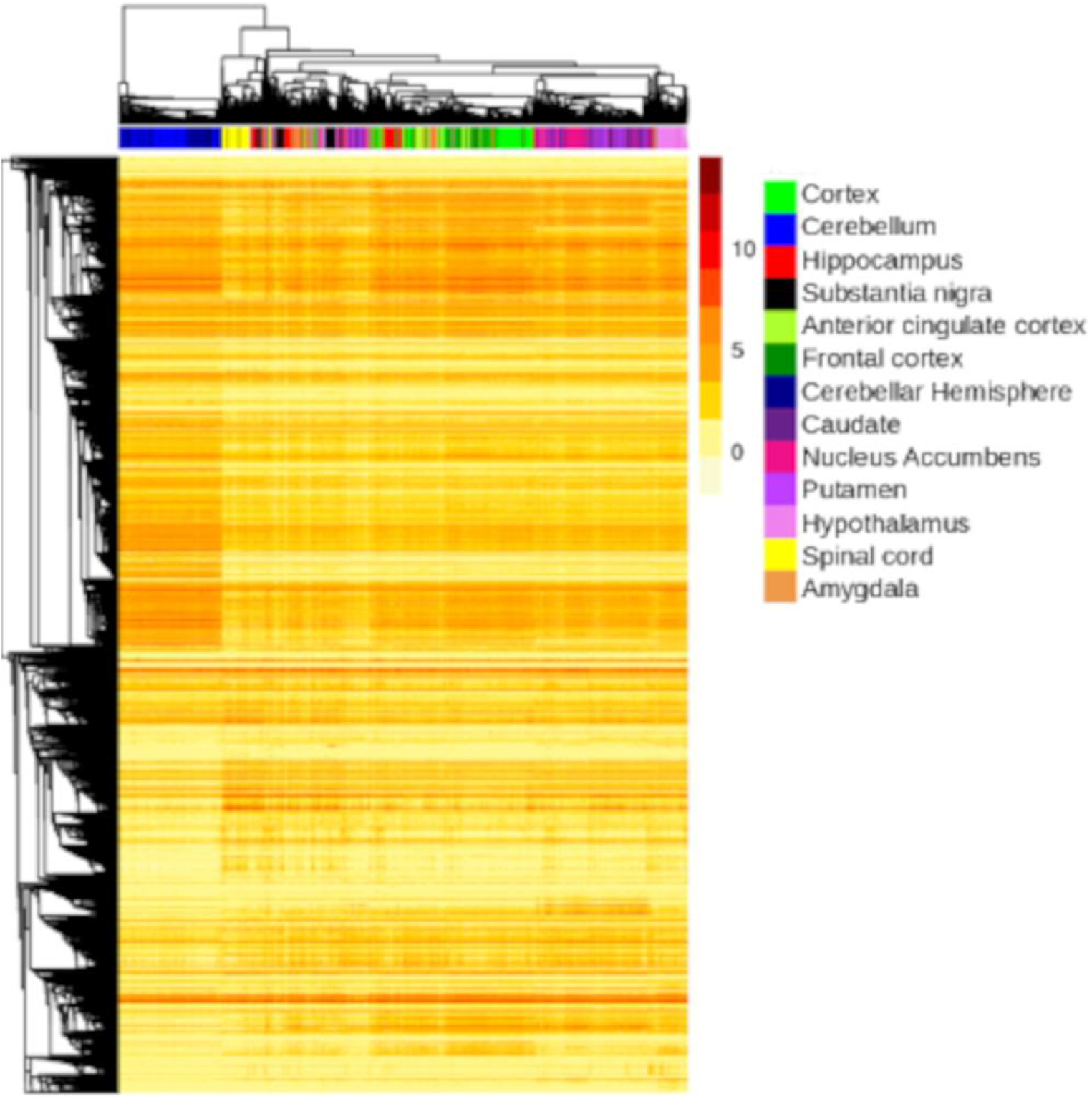
Unsupervised hierarchical clustering of genes expressed in human central nervous system regions. Hierarchical clustering of genes expressed in the CNS regions was performed based on gene pairwise distance in terms of correlation across gene expression values. The genes considered are paralogous genes. Each CNS region is represented by a different color. The regions belonging to the same anatomically defined CNS territories are represented in the same color: blue for the cerebellum region (cerebellum and cerebellar hemisphere regions), green for the cortex region (cortex, frontal cortex and anterior cingulate cortex regions), purple for the basal ganglia region (putamen, nucleus accumbens and caudate regions), and red for the amygdala-hippocampus region (amygdala and hippocampus regions). The remaining regions are considered as independent CNS regions: pink for the hypothalamus region, yellow for the spinal cord region and black for the substantia nigra.

In addition to this clustering analysis, we carried out another assessment by performing differential expression analysis of gene count data between all pairs of CNS regions (Methods). We obtained a list of significantly differentially expressed genes (DEGs) for each pair of regions (Additional File 2:Table S3). By comparing the relative proportion of DEGs in paralogs and singletons, we observed that DEGs were significantly enriched in paralogs for 75 out of the 78 region-pairs tested (Chi-squared test, and threshold p-value = 6.41E-04 with Bonferroni correction to account for the number of region pairs). Overall, these complementary expression studies using clustering and differential analysis illustrate the biological contribution of paralogous genes to expression differences between CNS territories. In Figure 1, the choice of color gradients for regions that anatomically overlap confirmed the ability of gene expression profiles to classify these regions into neurologically relevant groups. Therefore, from the next result sections on region-specificity analyses, we pool together some of the 13 initial regions that showed similar expression profiles in order to define a shorter list of 7 CNS regions (Methods).

### 2/ CNS region-specific expression of paralogs

We further investigated gene expression changes across the 7 CNS territories by looking at their region-specificity using the Tau score because of its high sensitivity to detect genes with narrow expression (25,26).

The Tau score ranges from 0 for broadly expressed genes, to 1 for highly specific genes (Methods). Contrary to Tau score distributions reported in a previous study of tissue-specificity on different organs (26), the distribution of Tau scores in the present study on intra-organ region-specificity was not bi-modal (Fig. 2A). Consequently, the Tau threshold for declaring a gene region-specific could not be visually defined. We thus developed an approach based on permutations to adapt this threshold choice to the case of regions within a single organ system. We calculated an empirical p-value for the Tau score of each gene, using permutations of the region labels to simulate the distribution of Tau scores in the absence of region-specificity, and then performed a False Discovery Rate (FDR) correction on the p-values for the multiple genes tested (Benjamini-Hochberg corrected p-value < 0.01) (Fig. 2A). This approach led to a Tau threshold of 0.525. We found that 17% (2,829) of protein-coding genes expressed in the CNS regions were region-specific. Moreover, we established that paralogs were significantly enriched in region-specific genes compared to singletons (19.2% of paralogs were region-specific, versus 13.9% of singletons, p-value = 2.045E-18, using a Chi-squared test) (Table 1). To check that low expression values did not bias the Tau score computation, we kept only genes with their maximal expression over the CNS regions higher than 1 RPKM and we obtained similar enrichment results (Additional File 2:Table S21). We confirmed this association between paralogs and region-specificity in addition to the effect of their expression level, by using a multivariate linear model that predicts the Tau score of a gene from its maximal expression over the CNS regions and its duplication status (Additional File 1:Result S1 and Additional File 2:Table S16A). This association was still observed when filtering out genes with low expression (<1 RPKM, Additional File 1:Result S1 and Additional File 2:Table S16A).

**Table 1.** Enrichments in CNS region-specific genes for the tested and reference gene groups

| Reference group<sup>a</sup> | Tested group for<br>CNS region-specificity<sup>a</sup> | Percentage of CNS |  |  |
| --- | --- | --- | --- | --- |
|  |  | region-specific genes<br>in the tested group<br>(%) | Chi-squared<br>test P-value<sup>b</sup> | Odds<br>ratio<sup>c</sup> |
| Protein coding<br>genes | Paralogous genes | 19.2 | 2.045E-18* | 1.48 |
| Paralogous genes<sup>d</sup> | WGD genes | 15.7 | 1.061E-18* | 0.64 |
|  | SSD genes | 22.6 | 9.022E-11* | 1.39 |
|  | ySSD genes | 28.6 | 6.341E-18* | 1.82 |
| SSD genes | ySSD genes | 28.6 | 3.483E-09* | 1.62 |
|  | oSSD genes | 15.6 | 2.729E-13* | 0.52 |
| WGD + wSSD genes | wSSD genes | 24.0 | 5.185E-12* | 1.69 |

<sup>a</sup> Abbreviations for gene duplication categories : WGD (Whole-Genome Duplication), SSD (Small-Scale Duplication), ySSD (younger SSD occurring after WGD events), oSSD (older SSD occurring before WGD events) and wSSD (WGD-old SSD occurring around WGD events).
<sup>b</sup> Application of Chi-squared tests (or of Fisher's exact test when the Chi-squared test could not be applied) with a corrected p-value threshold = 7.14E-03 (Bonferroni correction for 7 statistical tests).
<sup>c</sup> The odds ratio (>1 or <1) indicates the group (tested or non-tested respectively) in which there is an enrichment.
<sup>d</sup> The paralog reference group includes the genes belonging to WGD, SSD and WGD-SSD categories and the paralogs without annotation.

**Figure 2.**
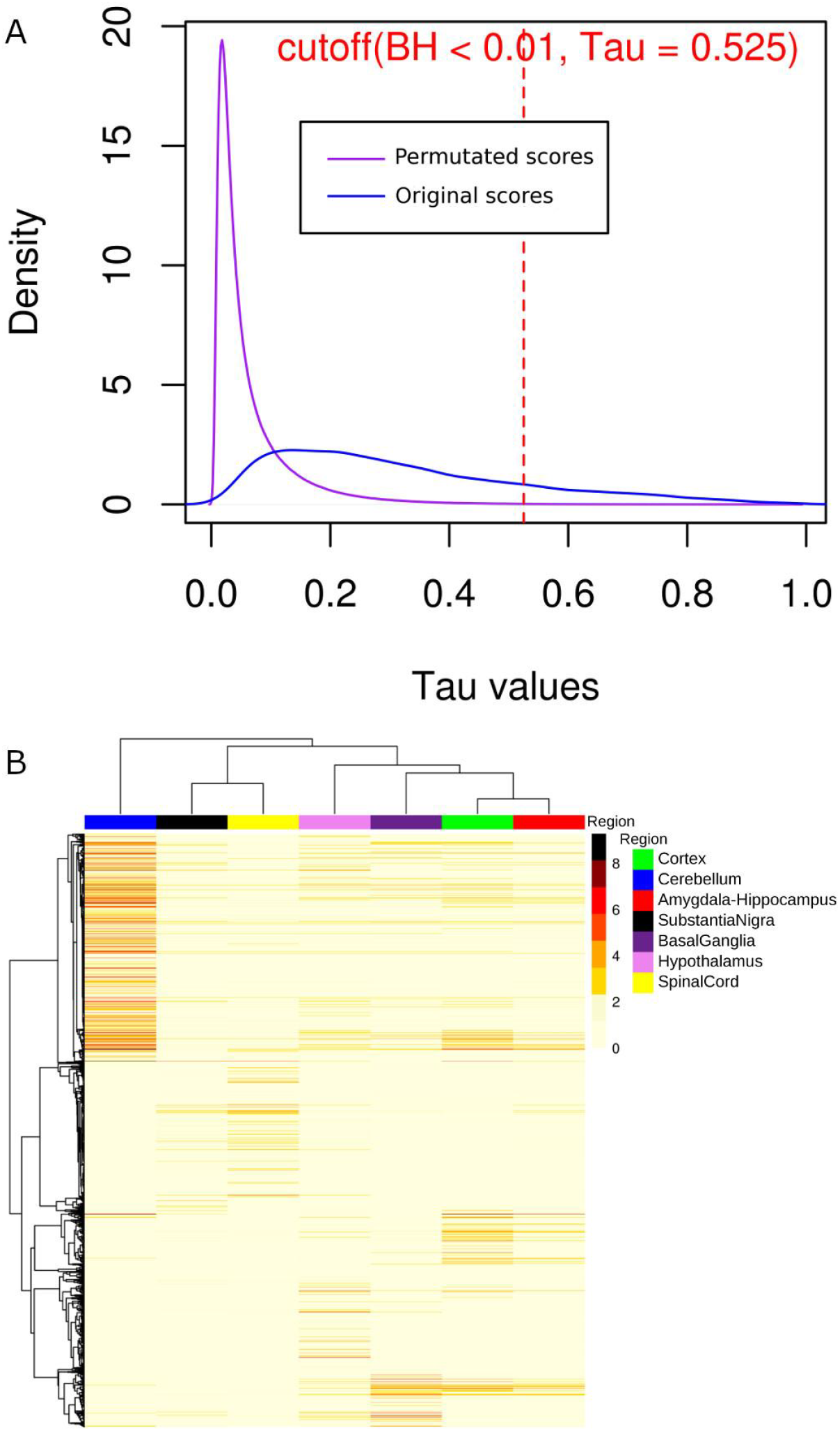
Specific expression of protein coding genes across human CNS regions. (A) Density plot of original Tau scores (blue line) calculated from the expression values of 16227 protein coding genes, and permutated Tau scores (purple line) calculated from 1000 x 16427 permutations. The region-specificity threshold of 0.525 (red dotted line) is defined, from permutated scores using the Benjamini-Hochberg corrected P-value of 0.01. (B) Unsupervised hierarchical clustering of region-specific genes expressed across CNS territories. The heatmap illustrates the mean gene expression calculated over samples of the cohort for each CNS region.

Although this method based on the Tau score can identify region-specific genes, it does not indicate which CNS region is targeted by this specificity (25). In order to study the regional distribution of gene expression, we mapped each specific gene to one CNS region (Additional File 2:Table S4). Therefore, for each region-specific gene, we considered the anatomical region associated with the highest expression value to be the specific region (Fig. 2B). We discovered that the distribution of region-specific genes across CNS territories was very heterogeneous (Additional File 2:Table S6) compared to an almost constant proportion of expressed genes across these regions (Additional File 2:Table S5). The highest proportions of region-specific genes were found in the cerebellum (40.2%), spinal cord (20.9%) and hypothalamus (16.4%). The remaining specific genes (22.5%) were scattered over the last four brain-related regions. The distribution of CNS region-specific paralogs was also highly heterogeneous and similar to the distribution obtained for all protein-coding genes (Additional File 2:Table S6).

In summary, we found that paralogs were more CNS region-specific than other genes. Furthermore, region-specific paralogs were concentrated in a limited number of CNS regions similarly to the other region-specific genes. Finally, we observed that beside the influence of abundance value, the paralog status also contributed to the specificity of gene expression to a CNS region.

### 3/ Evolutionary properties of CNS region-specific paralogs

The date of an SSD can be estimated in relation to the WGD events and attributed to one of the three duplication age categories: younger SSD (after WGD events - ySSD), older SSD (before WGD events-oSSD) and WGD-old SSD (around WGD events – wSSD) (Methods) (27). Using our collection of paralogs with CNS region-specific expression, we performed statistical tests to determine if they were enriched in particular duplication events (WGD or SSD) or dates of SSDs (oSSD, wSSD and ySSD categories). Genes can undergo both WGD and SSD duplication and can sometimes be retained after each duplication. Unless otherwise stated, when we refer to a duplication type from this point on in the paper, we are referring to genes that have been retained after this duplication type only (WGD or SSD), in order to make a clear distinction between the effects of the two duplication types. Of the 10,335 paralogs considered in our study, 5,114 are from WGD, 3,719 from SSD (1,192 from ySSD, 1,260 from wSSD and 1,267 from oSSD) and 1,502 unclassified (966 both WGD-SSD and 536 without annotation). We first observed that among paralogs, SSD genes were significantly enriched in CNS region-specific genes (22.6% of SSDs were region-specific versus 17.3% of the other paralogs, p-value = 9.022E-11), while on the opposite WGDs were depleted in region-specific genes (Table 1). Furthermore, when we performed the same analysis only on the paralogs duplicated around the WGD events (WGDs and wSSDs) to remove the potential confounding effect of the duplication date, the WGD genes were still significantly depleted in region-specific genes (15.7% of WGDs were region-specific versus 24% of wSSDs, p-value = 5.185E-12) (Table 1). This last test allowed us to conclude that SSD paralogs were enriched in CNS region-specific genes, independently of the potential effect of the duplication date.

In addition to assessing the effect of duplication type, we also tested the association between duplication age categories and region-specificity, and found that ySSD were even more enriched in region-specific paralogs (28.6% of ySSDs versus 18.0% of the remaining paralogs, p-value = 6.341E-18). This last result was confirmed by the fact that ySSDs were still enriched in region-specific paralogs when we performed the analysis on SSD paralogs only (28.6% of ySSDs versus 19.8% of the remaining SSDs, p-value = 3.483E-09). On the other hand, oSSDs were depleted in region-specific genes compared to other SSD paralogs (15.6% of oSSDs versus 26.2% of the remaining SSDs, p-value = 2.729E-13) and showed the same proportion of region-specific genes as WGDs (15.7%) (Table 1).

Expression of young duplicates has been evidenced to be lower than older duplications (Guschanski et al, 2017). Thus, we have further explored the expression levels of the different types of genes (Singletons, WGDs, oSSDs, wSSDs and ySSDs) and their influence on region-specificity. First, we reported the distribution of singletons and the different duplicate types through bins of expression values (Figure 3A). While the singletons and the WGD and oSSD duplicates were distributed among the expression bins according to a Gaussian profile with a peak for the range of 7 to 15 RPKM, the distribution of ySSDs had its maximum for low levels of expression and then decreased progressively towards the highest abundances. This result confirmed that ySSDs tended to be more weakly expressed within the CNS than other types of genes. We then questioned whether the low levels of expression were associated with a higher CNS region-specificity for the different types of genes (Figure 3B). The distribution of the proportions of region-specific genes per bin of expression showed that approximately 50% of genes expressed in the range of 0 to 1 RPKM were region-specific, whatever the type of genes. On average, these proportions decrease with increasing expression levels up to 63 RPKM. More precisely, we observed for the expression bins in the range of 1 to 31 RPKM, greater proportions of region-specific genes for the wSSD and ySSD types, compared to the other gene types. In addition, an increase in the proportion of region-specific genes for the oSSD and wSSD types seems to appear in the range 63 - 127 RPKM. However, we noted that for higher expression bins (> 127 RPKM) the number of genes was not enough in some gene types to compare their proportion of region-specific genes. Finally, the gradual decrease in the percentage of regions-specific genes with increasing levels of expression suggests that this trend is more related to a biological reality than a technical effect. However, we cannot completely rule out the possibility that the calculation of the Tau score could be biased for genes characterized by low expression values. Therefore to assess this potential bias, we performed the same enrichment tests as in Table 1 by removing weakly expressed genes (genes with their maximal expression over the CNS regions lower than 1 RPKM), and we confirmed the enrichment of SSDs and particularly of ySSDs in region-specific genes (Additional File 2:Table S21).

**Figure 3.**
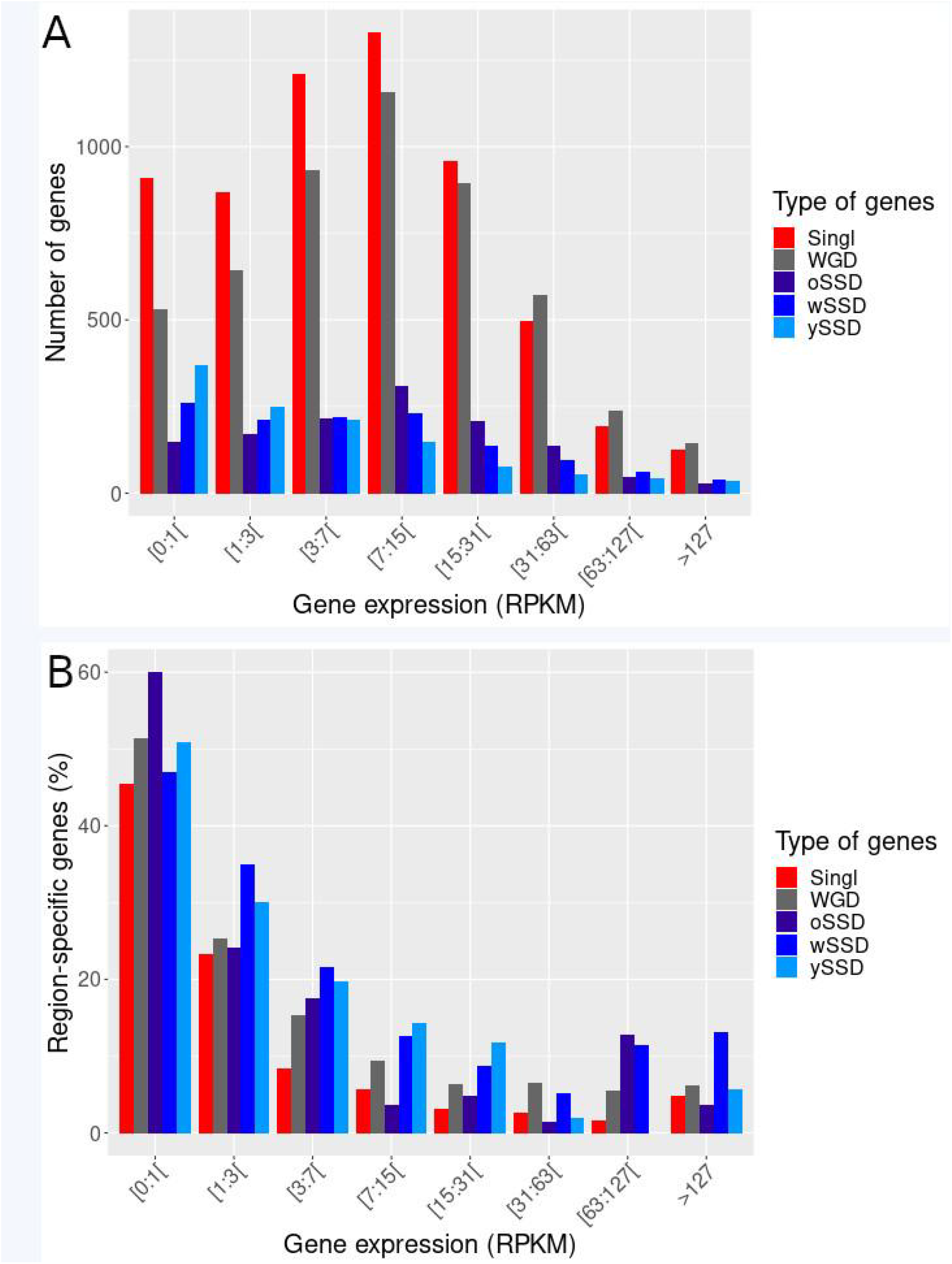
Distribution of CNS region-specific genes across ranges of expression values. Barplots show (A) the number of expressed genes and (B) the percentage of region-specific genes for different expression bins. For each gene, we first calculated its expression value per CNS region by averaging over all the samples associated with each region. We then selected as reference value for each gene, the maximum of these averages of expression across the CNS regions. Gene expression values are given in RPKM (on a log2 scale) and each bin corresponds to 1 unit of the log2(RPKM + 1) values. The last bin groups all gene expressions higher than 255 RPKM.

We also confirmed the contribution of both duplication age and duplication type to the region-specificity of paralogs, independently of the effect of their expression level, using multivariate linear models (Additional File 1:Result S1 and Additional File 2:Tables S16B-C).

To obtain a complementary view of this region-specificity for recent duplications, we examined the distribution of the Tau scores of paralogs according to their phyletic age (Fig. 4). We found that the maximum Tau scores were obtained for genes with phyletic ages around 0.12 which corresponds in most cases to ySSD duplication events that occurred around the separation of the Simians clade (Ensembl Compara GRCh37 p.13).

**Figure 4.**
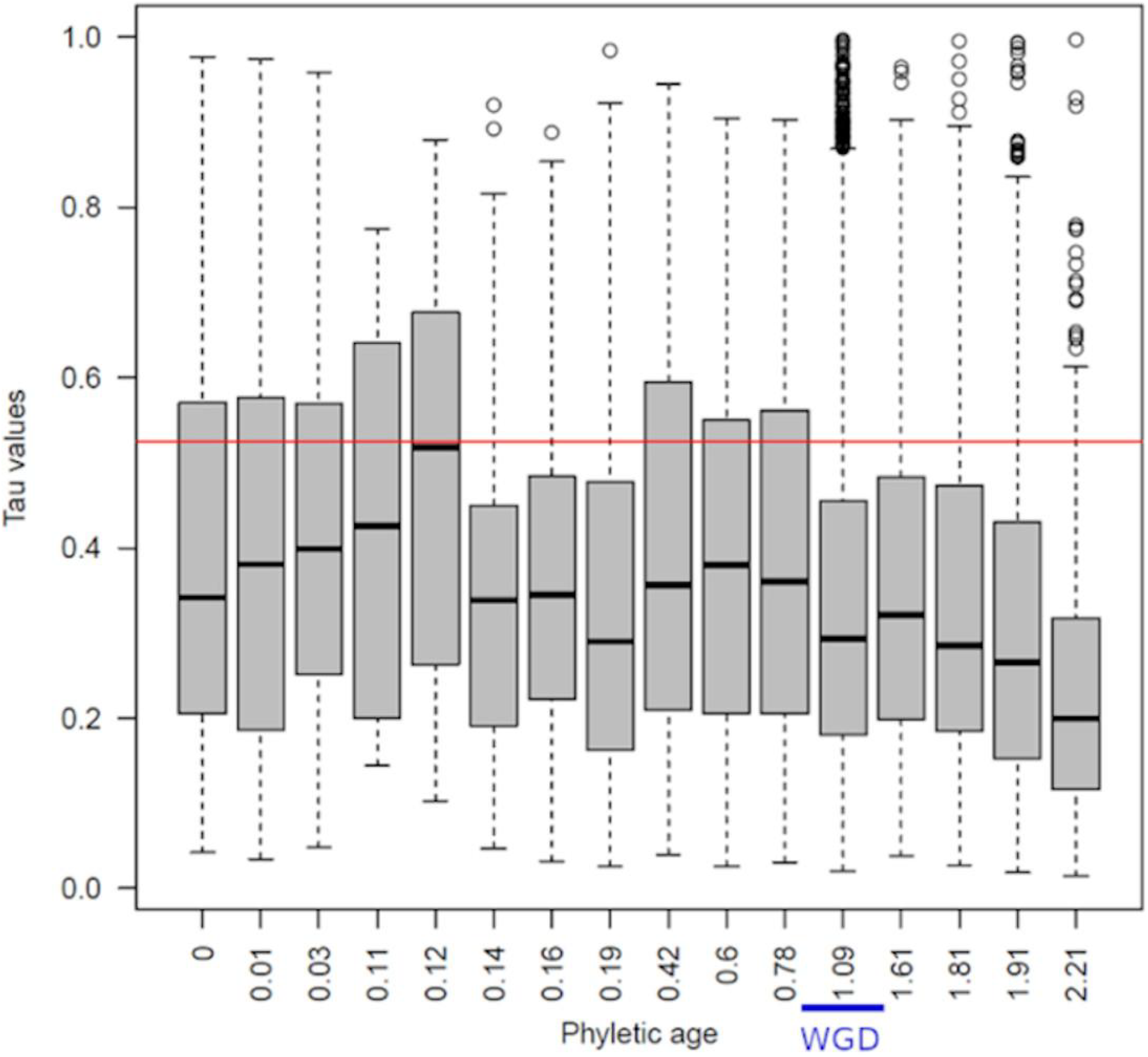
Association between the phyletic age of the duplication and the region-specificity. Boxplots show the distribution of Tau scores for paralogs grouped according to their phyletic age obtained from Chen et al., 2013. The range of phyletic ages corresponding to WGDs is indicated by a blue horizontal bar. The red horizontal line represents the threshold of region-specificity (Tau score = 0.525).

In summary, we found that SSD genes and in particular ySSD genes were more CNS region-specific than other paralogs, probably due to both their SSD origin and their duplication age, in addition to the influence of expression level on region-specificity.

### 4/ CNS region-specific expression of gene families

We previously found that paralogs, and especially SSDs and ySSDs, were involved in territorial expression of CNS regions. Paralogs being organized into gene families, we also assessed whether or not the paralogs belonging to the same family tend to share region-specific expression properties.

We first studied the expression similarity between paralogs across CNS regions by using a co-expression analysis without using *a priori* knowledge on their region-specificity. The study of co-expression allowed us to explore the higher level of organization of the paralogs into groups of genes with coordinated expression across CNS regions and compare these modules of co-expressed paralogs across regions against annotated gene families. The Weighted Gene Correlation Network Analysis (WGCNA) methodology (28) was used to infer the correlation-based co-expression network. Contrary to previous studies that inferred a network per tissue and then compared modules between networks (29,30), we carried out co-expression network inference by simultaneously using all the 13 CNS region samples profiled by the GTEx consortium in order to explore gene associations with region differentiation. We optimized the WGCNA to generate highly correlated co-expression modules of small size in order to compare them with the annotated gene families (Methods and Additional File 1:Figure. S3). Indeed, out of our 3,487 gene families, 1,644 (47%) were constituted of only two genes. Our WGCNA analysis extracted 932 modules of co-expressed paralogous genes. Only 104 genes were not included in a co-expression module. The module size ranged from 2 to 911 genes with 84% of small size modules (modules with less than 10 genes) (Additional File 2:Table S7). If we consider modules greater than 20 genes, a high proportion of them were enriched in molecular function and biological process GO terms indicating that our network inference approach captured shared biological functions among co-expressed paralogs (Additional File 1:Result S4).

This co-expression network analysis allowed us to classify the gene families into two categories, homogeneous and heterogeneous gene families, based on their patterns of expression across CNS regions (Methods). A homogeneous gene family was defined by the property that the majority of its member genes were included in the same co-expression module. Out of the 3,487 gene families considered in this co-expression study, we identified 111 homogeneous families (with 257 co-expressed paralogs out of a total of 300 expressed paralogs in these families, the remaining 43 not co-expressed paralogs being removed from all tests on homogeneous family genes in the rest of the article) and thus 3,376 heterogeneous families (10,035 paralogs) (Additional File 2:Tables S13-S14). We showed by a permutation approach that this number of homogeneous families was significantly large, with an empirical p-value inferior to 10^−3^ (Methods), suggesting that paralogs were more co-expressed across CNS regions when they came from the same family. A biological pathway enrichment analysis of the homogeneous family genes revealed that they were notably enriched in transcription factors and signaling proteins involved in neural development (Additional File 1:Result S6 and Additional File 2:Table S10).

We then investigated the link between shared region-specificity and homogeneous gene families by categorizing families according to their region-specificity (31) (Additional File 1:ResultS7). Families composed of a majority of genes specific to the same regions were classified as region-specific families. We identified 58 region-specific families and we found a significant enrichment of these families in homogeneous families (45% of region-specific families versus 2.5% of other families, p-value = 1.691E-69) (Table 2).

**Table 2.**
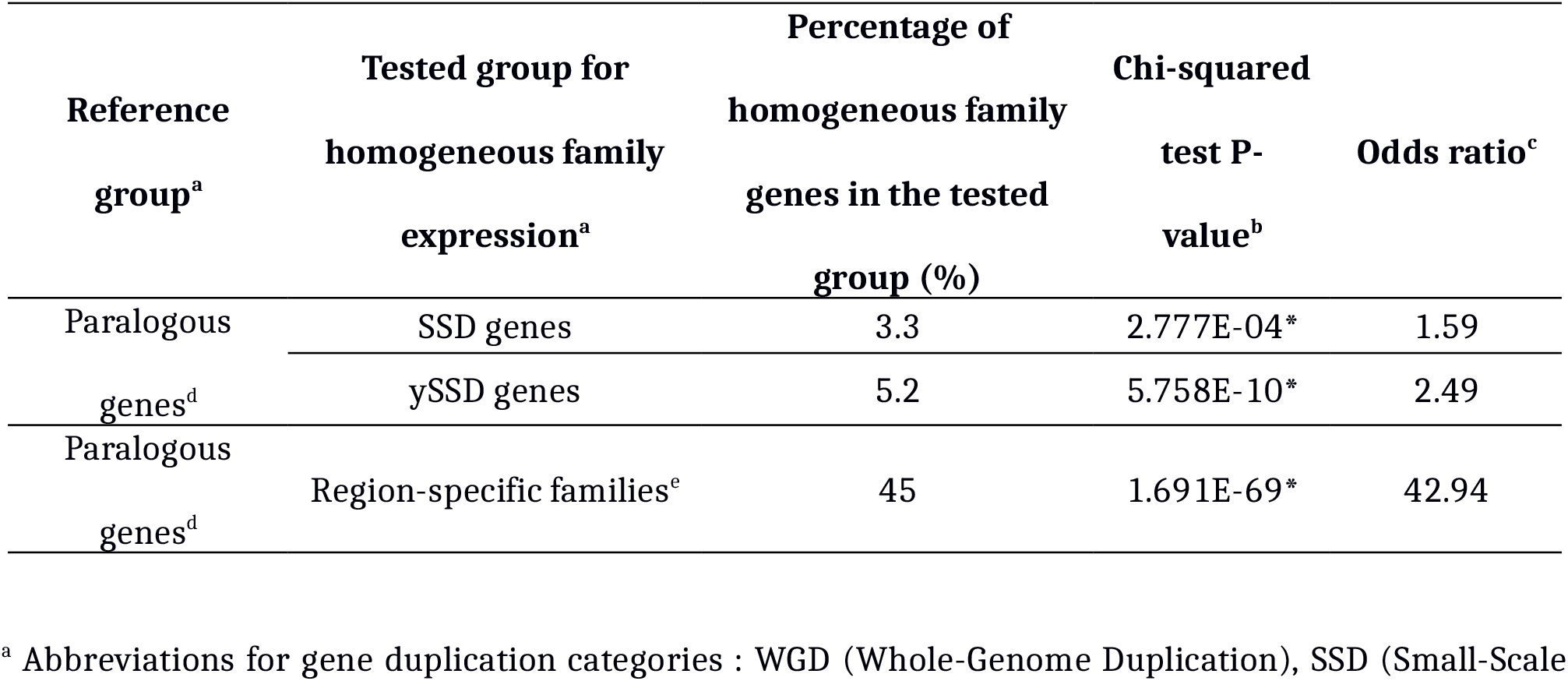
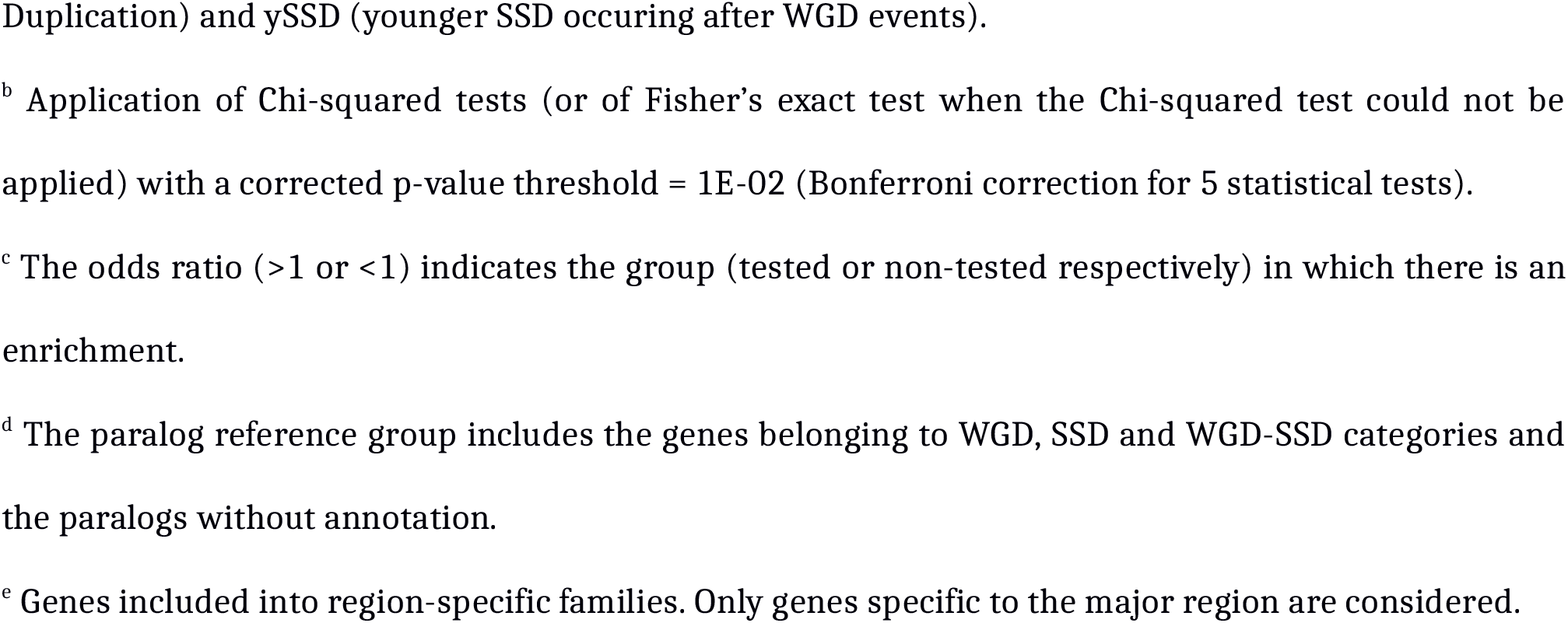
Enrichments in genes from homogeneously expressed families for the tested and reference gene groups.

| Reference<br>group<sup>a</sup> | Tested group for<br>homogeneous family<br>expression<sup>a</sup> | Percentage of<br>homogeneous family<br>genes in the tested<br>group (%) | Chi-squared |  |
| --- | --- | --- | --- | --- |
|  |  |  | test P-value<sup>b</sup> | Odds ratio<sup>c</sup> |
| Paralogous<br>genes<sup>d</sup> | SSD genes | 3.3 | 2.777E-04* | 1.59 |
|  | ySSD genes | 5.2 | 5.758E-10* | 2.49 |
| Paralogous<br>genes<sup>d</sup> | Region-specific families<sup>e</sup> | 45 | 1.691E-69* | 42.94 |

<sup>a</sup> Abbreviations for gene duplication categories : WGD (Whole-Genome Duplication), SSD (Small-Scale
Duplication) and ySSD (younger SSD occurring after WGD events).
<sup>b</sup> Application of Chi-squared tests (or of Fisher's exact test when the Chi-squared test could not be applied) with a corrected p-value threshold = $1E-02$ (Bonferroni correction for 5 statistical tests).
<sup>c</sup> The odds ratio ( $>1$ or $<1$ ) indicates the group (tested or non-tested respectively) in which there is an enrichment.
<sup>d</sup> The paralog reference group includes the genes belonging to WGD, SSD and WGD-SSD categories and the paralogs without annotation.
<sup>e</sup> Genes included into region-specific families. Only genes specific to the major region are considered.

Finally, we studied whether homogeneous families were associated with a type of duplication event or with a duplication age. We found that SSD genes and ySSD genes in particular were enriched in genes coming from homogeneous families (3.3% of SSD versus 2.1% of the other paralogs, p-value= 2.777E-04; 5.2% of ySSD versus 2.1% of the other paralogs, p-value = 5.758E-10) (Table 2). Similarly, SSD and ySSD genes were significantly enriched in genes coming from region-specific families (Additional File 2:Table S17). Lastly, we also investigated the shared region-specificity at the paralog pair level. We observed, among region-specific pairs, a high proportion of SSD (50%) and ySSD (59%) pairs specific to the same region than WGD pairs (31%), even though the very low number of these region-specific pairs did not allow us to get significant results (Additional File 1:Result S2).

In order to interpret these results, one may expect that the co-expression level between two duplicates in a paralog pair will be associated with their proximity on the genome, as epigenetic co-regulation of gene expression partly depends on the proximity between genes on the genome (32–34). We thus investigated whether the genomic distance between paralog pairs (Additional File 1:Result S5) could be used to differentiate homogeneous from heterogeneous families. For homogeneous families, we considered only pairs in which both paralogs belonged to the main co-expression module (37 pairs), and removed the other pairs from the test. We found that homogeneous families were depleted in inter-chromosomal pairs (70.3% of homogeneous families versus 90.2% of heterogeneous families were spread across different chromosomes, p-value= 7.73E-04) and were enriched in tandem duplicated pairs (27% of homogeneous families and 6.7% of heterogeneous families were separated by less than 1 Mb, p-value = 1.743E-04) (Additional File 2:Table S20); this supports the idea that paralog co-expression is favored by proximity along the genome. Moreover, we confirmed that the genomic proximity of duplicates was associated with recent SSDs and that the younger the SSD pair, the more the duplicates were found in tandem in the genome (Additional File 1:Result S5). The tandem duplication explains why SSDs, and especially ySSDs, tend to be more co-expressed and to share more often the same region-specificity within their family than other paralogs.

In summary, the gene co-expression network analysis performed on the CNS regions allowed us to find that within gene families, the shared region-specificity of paralogs was associated with their co-expression across regions and we classified gene families into two categories according to co-expression status. Homogeneous families were enriched in paralog pairs which were closely located on the genome in tandem duplication, probably due to the specific trend of SSD pairs to be duplicated in tandem. Indeed, these homogeneous families were enriched in SSDs, especially in ySSDs, and were associated with a shared region-specificity.

## DISCUSSION

As far as we are aware, this study is the first to focus specifically on the spatial expression of paralogs and gene families between the different human CNS territories based on post-mortem human tissues analyzed by the GTEx consortium. Previous studies based on gene expression analysis between organs have already established the important association between paralogs and tissue differentiation (10,35). We showed the contribution of paralogs to expression differences between CNS territories.

Paralogs are known to be more tissue-specific than other genes (10,31,36,37). Among paralogs, SSDs (15) and in particular ySSDs (35) seem to be more often tissue-specific than other paralogs when comparing tissues from different organs. However, when considering the brain as a whole and comparing it with other organs, it has been found that WGDs tend to be enriched in brain-specific genes compared to SSDs (15,16,31). In order to obtain a complementary vision to these previous studies, we focused on the expression of paralogs considering only the regions that composed the human CNS. We observed that paralogs, especially ySSDs were more region-specific than other genes. In addition, we found that even wSSDs were enriched in region-specific genes compared to other paralogs of the same age (WGDs), thus suggesting that the region-specificity between brain regions is not only associated with the young age of duplication but also with the type of duplication (i.e. with SSD duplications). Our results, although apparently contradictory, do not question the known involvement of WGDs in brain-specific expression. Indeed the genes specific to the brain as a whole may not be specific to a particular CNS region and conversely a gene specific to a given region within the brain may also be expressed in other organs. Moreover, the fact that an SSD gene tends to be more often specific to only one or just a few CNS anatomical regions than a WGD gene, implies that the average expression of SSD genes over the whole brain would be lower than the average expression of WGDs. Thus, this broad expression of WGDs within the brain regions facilitates the detection of their brain-specific expression when comparing several organs, while some ySSDs specific to the brain may not be detected.

Using multivariate linear models, we reported the major contribution of expression level and that of duplication status to region-specificity in CNS territories. Among paralogs, we found that the SSD duplication type explained also part of the region-specificity variance. Regarding the evolutionary time, low phyletic ages were also significantly associated with high region-specificity; a property potentially restricted to CNS regions. Beside this global effect of the duplication age, we observed that the highest region-specificity seemed to occur for young duplication events, around the separation of the Simians clade.

We then studied the gene family level of organization using gene co-expression network analysis of paralogs across CNS regions. We showed that modules of co-expressed genes were able to identify clusters of paralogs with the same region-specificity. The characterization of gene families according to the level of co-expression of their member genes has led to the identification of two categories of families: homogeneous families, which are composed of a majority of co-expressed genes, and heterogeneous families. We observed that homogeneous families were enriched in ySSD genes and tandem duplicate pairs, in agreement with a previous study showing that pairs of ySSD paralogous genes tend to be duplicated in tandem and co-expressed just after the duplication event (11). A previous study established that when the two paralogs of an ySSD pair are tissue-specific, they tend to be specific to the same tissue more often than for other paralog pairs (35). Similarly, regarding region-specificity in the CNS, we showed the high co-expression of ySSD pairs and the enrichment of co-expressed families in region-specific families, where the majority of genes were region-specific to the same region.

From the analysis of gene expression across human and mouse organs, Lan and Pritchard 2016 proposed a model for the retention of SSD duplicates appearing in mammals. In this model, pairs of young paralogs are often highly co-expressed probably because tandem duplicates are co-regulated by shared regulatory regions. In addition, this model is consistent with the dosage-sharing hypothesis in which down regulation of the duplicates, to match expression of the ancestral gene, is the first step enabling the initial survival of young duplicates (11). Our analyses of ySSDs expression features between CNS territories seem to be concordant with this model, indeed ySSDs tend to be organized within small families of co-expressed genes and also weakly expressed in concordance with the sharing of the gene ancestral expression. Furthermore, our results in the CNS regions seem to confirm that, after the initially high co-expression of SSD paralogs just after their duplication, they become more region-specific and less co-expressed in part through chromosomal rearrangement, suggesting a long term survival by sub-/neofunctionalization (11).

## CONCLUSIONS

Our exploration of paralogs suggests that young SSDs are particularly involved in the specificities of expression of the different human CNS territories. This suggests the relevance to investigate paralog expression between the territories of the same organ. However, to determine whether or not the region-specific expression patterns of young SSDs are solely associated with the central nervous system regions, it will be interesting to explore their expression between anatomic regions of other complex organs.

## METHODS

### Human genes, duplication events and families

A list of 21,731 human genes, with both their HGNC gene symbol and their Ensembl IDs (GRCh37, release 59), was collected based on the work of Chen and co-workers (38). Among these genes, 14,084 paralogs made up of 3,692 gene families, identified by TreeFam methodology (39), were obtained from Chen and co-workers (38). These authors downloaded all gene families from the TreeFam v.8.0 database, which identifies duplicates based on gene family evolution. Moreover, for each paralog, they represented the phyletic age of its last duplication event by the total branch length from the node indicating where the duplication event had happened on the species tree to the human leaf node, and they assigned the associated duplicate (38,40). A second list of 20,415 genes was extracted from Singh *et al.* 2014. This gene ID list was converted to HGNC gene symbols and intersected with the first list in order to annotate it (17,805 protein-coding genes in common). Thus, in the present study, we collected the duplication category for each paralog (27) (Singh et al. 2014). Singh et al. obtained WGD annotations from (41) and obtained their SSD annotations by running an all-against-all BLASTp using human proteins (42). Singh and co-workers defined genes as singletons if they were not classified as WGDs or SSDs and they obtained the duplication age for SSD genes from the Ensembl compara (43). They classified paralogs into the following categories: WGD, SSD, ySSD (i.e. SSD with duplication event younger than WGD), oSSD (i.e. SSD with duplication event older than WGD) and wSSD (i.e. SSD with duplication date around the WGD events). There were 5,390 annotated paralogs originating from the WGD and 4,889 from SSD (2,104 from ySSD, 1,354 from oSSD and 1,431 from wSSD). Moreover, there were 2,607 paralogs without annotations and 1,198 paralogs annotated as both WGD and SSD (WGD-SSD). The WGD-SSD paralogs were not included into the WGD or the SSD duplication categories. However, the unannotated and WGD-SSD paralogs were both considered into the paralog group. We verified that these paralog duplication categories were consistent with the phyletic ages (duplication dates) collected from Chen and co-workers (38,40) (Additional File 1:Figure S4). The list of our paralogous gene pairs and families is given in the supplementary table S1 (Additional File 2:Table S1). The evolutionary annotation of paralogous genes is indicated in the supplementary table S2 (Additional File 2:Table S2). The list of singleton genes is given in the supplementary table S12 (Additional File 2:Table S12). Furthermore for the analysis of the duplicate pairs, we considered only the 3,050 pairs which appeared twice in our paralog list (i.e. where the first paralog is associated with the second paralog and vice versa and where the duplication category annotation is the same for both paralogs); genomic distances between duplicate pairs were obtained from Ensembl (GRCh37/90).

### Gene expression profiles in CNS regions

We obtained gene counts and RPKM (Reads Per Kilobase Million) values for 63 to 125 individuals (1259 post-mortem samples – RNA integrity > 6) distributed over 13 CNS regions(cerebellum, cerebellar hemisphere, cortex, frontal cortex, anterior cingulate cortex, hypothalamus, hippocampus, spinal cord, amygdala, putamen, caudate, nucleus accumbens and substantia nigra) from the GTEx consortium data release 6 (GRCh37) (24). The CNS regions associated with each GTEx patient sample used in our study is indicated in the supplementary table S11 (Additional File 2:Table S11). These gene expression data, calculated by GTEx took into account only uniquely mapped reads (https://gtexportal.org). We filtered out low-information content genes by removing genes with a null variance across samples and weakly expressed genes with mean expression per region lower than 0.1 RPKM for all regions. We thus kept for analyses a total 16,427 genes distributed across 10,335 paralogs (5,114 WGD, 3,719 SSD, 1,192 ySSD, 1,260 wSSD and 1,267 oSSD, 966 WGD-SSD and 536 without annotations) grouped in 3,487 families and 6,092 singletons. It should be noted that all our analyses were performed on this list of expressed genes only. Gene RPKM values were log-transformed (log2 (RPKM + 1)) and adjusted by linear regression for batch effects and various biological effects (sequencing platform, age, gender and the first 3 principal components of genetic data illustrating the population structure given by the GTEx Consortium; the intercept of the regression was not removed from the residuals in order to keep the mean differences between genes (https://www.cnrgh.fr/genodata/BRAIN_paralog)). These filtered, log-transformed and adjusted RPKM values were used as input for unsupervised classification of brain regions, as well as for gene co-expression network inference and for region-specificity analysis. Moreover, gene expression data for regions considered to anatomically overlap were merged by calculating the average expression value across related regions prior to the expression specificity analysis. Therefore, from an initial list of 13 regions, we gathered samples into a shorter list of 7 CNS regions: cerebellum (cerebellum and cerebellar hemisphere), cortex (cortex, frontal cortex and anterior cingulate cortex), basal ganglia (putamen, nucleus accumbens and caudate), amygdala-hippocampus, hypothalamus, spinal cord and substantia nigra.

### Unsupervised clustering of gene expression profiles

Gene expression profiles (filtered and adjusted RPKM values) generated by the GTEx Consortium for the 1,259 samples distributed across the 13 CNS regions, were clustered by unsupervised hierarchical clustering using the pheatmap package of R version 3.4 (similarity measure: Pearson correlation, clustering method: average linkage).

### Differential gene expression analysis

Genes with low-information content were removed before differential gene expression (DGE) analysis. DGE analysis was performed by DESeq2 (44) on count data for each pair of CNS regions, with the “median ratio” between-sample normalization and using batch and biological effects as covariates. For each region pair, we then corrected gene p-values for the number of tested genes using FDR (45) and obtained a list of significantly differentially expressed genes (DEGs) (FDR<0.05). Finally, we considered only the DEGs with an absolute log2 fold-change greater than 0.5.

### CNS region-specificity calculation

#### Tau score calculation

To identify genes expressed in specific regions of the CNS, we used the τ score that was proposed to estimate the degree of tissue-specificity of each gene (25) :

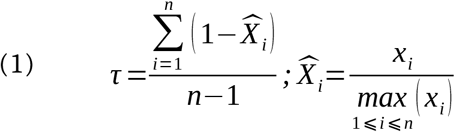

In this equation, *x*_*i*_ is the mean expression of a given gene in region *i* and *n* is the number of different regions. τ varies from 0 to 1 where 0 indicates that the gene is broadly expressed and 1 that the gene is tissue-specific. For τ computation, genes must have a positive mean of expression in every CNS region. Although we log-normalized expression data with log2(RPKM+1) leading to positive expression values, the correction for batch and some biological effects induced some negative values in gene mean expression. We pooled expression data generated by GTEx for the 13 regions into the list of 7 non-overlapping CNS regions so that the τ score would not decrease artificially for genes specific to several close sub-regions.

#### Tau score threshold defined by permutations

The τ score was computed for each gene and for the 7 CNS regions. We then plotted the τ score distribution obtained from all protein coding genes (Fig. 2A). However, there is no general τ score threshold at which a gene is considered to be region-specific. To define a region-specificity threshold, we implemented a statistical method based on permutations. We applied 1000 permutations on the region labels assigned to the samples to shuffle the correspondence between samples and regions. For each permutation, τ scores were recomputed for each gene. The distribution of the 1000 × 16427 τ scores obtained from the permutations is given in Figure 2. For each gene and its original τ score, a p-value was then calculated as the proportion of permutation-based τ scores higher than the original τ score. The Benjamini-Hochberg correction for the number of genes tested was applied to all p-values. Genes with a corrected p-value lower than 0.01 were declared CNS region-specific, which corresponded to a τ score threshold of 0.525 (Fig. 2A). Visualization of gene profiles across brain regions at different windows of the τ score showed region-specificity beyond the τ score threshold of 0.525. We visualized expression values at different windows of Tau scores and we observed better region-specific profiles over 0.5 (Additional File 1:Figure. S2). Therefore for each region-specific gene, we considered that the CNS region with the highest expression value to be the specific region.

### Inference of gene co-expression networks

The gene network inference was carried out using the Weighted Gene Correlation Network Analysis (WGCNA) methodology (28), which generates co-expression networks and identifies modules (groups) of co-expressed genes. We applied the WGCNA tool only to paralogous gene expression data (RPKM) across the GTEx samples of the 13 CNS regions. Genes were grouped into modules according to their expression profile similarity. The module named “grey”, which grouped genes that were not considered as co-expressed by WGCNA, was composed of genes with very low variability across all samples. Since we had removed the genes with no variance across region samples and those which were weakly expressed before performing the WGCNA analysis, the grey module was small in size (104 genes). Furthermore, if this filtering had not been performed, some of the genes with an overall weak expression might have been integrated into co-expression modules, thus creating a bias. One of our goals was to compare gene families to co-expression modules. Given that 47% of gene families have a size equal to 2, we optimized WGCNA parameters to obtain small highly co-expressed modules (Additional File 1:Result S3).

### Homogeneous and heterogeneous families

#### Definition

A gene family was defined as homogeneous if the majority, more than 60%, of its member genes were included in the same co-expression module. It should be noted that the total size of gene families was used to compute this percentage, even if some member genes were not in the list of expressed paralogs. Gene families which did not respect this homogeneity rule, i.e. those with member genes scattered over different co-expression modules, were defined as heterogeneous.

#### Assessment of the significance of the number of homogeneous families

Starting from the paralog modules obtained with WGCNA, we used a permutation procedure (by permuting 1,000 times the module labels of paralogs and counting the number of falsely homogeneous families for each permutation) and were able to conclude that the number of homogeneous families was significantly large, since for each permutation the number of falsely homogeneous families was lower than the number that we obtained, leading to an empirical p-value inferior to 10^−3^.

## LIST OF ABBREVIATIONS

CNS: Central Nervous System
DGE: Differential Gene Expression
DEG: Differentially Expressed Gene
FDR: False Discovery Rate
GTEx: Genotype Tissue-Expression
oSSD: SSD with duplication date older than WGD events
RPKM: Reads Per Kilobase per Million mapped reads
SSD: Small Scale Duplication
WGCNA: Weighted Gene Co-expression Network Analysis
WGD: Whole Genome Duplication
wSSD: SSD with duplication date around the WGD events
ySSD: SSD with duplication date younger than WGD events

## ETHICS APPROVAL AND CONSENT TO PARTICIPATE

Not applicable

## CONSENT FOR PUBLICATION

Not applicable

## AVAILABILITY OF DATA AND MATERIALS

Raw data analyzed in our study can be found at https://gtexportal.org and processed data can be found at https://www.cnrgh.fr/genodata/BRAIN_paralog.

All data generated or analysed during this study are included in this published article and its supplementary information files.

## AUTHORS CONTRIBUTIONS

SBJ, VF, ELF and CB conceptualized the project and the methodology. They also interpreted the results and wrote the paper. The computational and statistical analyses, and their visualizations were done by SBJ, SC, ELF and CB. VF, ELF and CB supervised the study. VF, JFD, ELF and CB participated in the funding acquisition. VM and JFD reviewed the paper. All authors read and approved the final manuscript.

## ACKNOWLEDGMENTS

We are grateful to Marc Robinson-Rechavi for his feedback on the methods, the results and the result interpretation. We thank Steven McGinn and Elizabeth May for English language editing. We also thank Carène Rizzon, Margot Coréa, Olivier Jaillon, François Artiguenave and Morgane Pierre-Jean for constructive discussions.

## ADDITIONAL FILES

### Additional File 1 (PDF format)

**Result S1.** Multivariate linear regression models to explain the Tau score.

**Result S2.** Association between region-specific expression in the same region and paralog pairs.

**Result S3.** Optimization of WGCNA parameters.

**Result S4.** Biological relevance of co-expression modules.

**Result S5.** Genomic distances between pairs of paralogs.

**Result S6.** Characterization of homogeneous and heterogeneous gene families.

**Result S7.** Association between co-expression and shared region-specificity.

**Figure S1.** Unsupervised hierarchical clustering of genes expressed in human central nervous system regions.

**Figure S2.** Unsupervised hierarchical clustering of region-specific gene expression across CNS regions for different Tau score intervals.

**Figure S3.** Optimization of Weighted Gene Co-expression Network Analysis (WGCNA) parameters.

**Figure S4.** Comparison between gene duplication dates generated by (Chen et al. 2013) and by (Singh et al. 2014).

**Figure S5.** Comparison between original and permuted Tau scores of protein coding genes across human CNS regions (Expression threshold > 1 RPKM).

### Additional File 2 (XLSX format)

**Table S1.** Paralogous gene pairs and gene families.

This table describes the evolutionary information collected for each pair of duplicated protein-coding genes in humans.

<u>HUGO_geneSymbol</u> : Homo sapiens gene name based on HUGO nomenclature

<u>ENSEMBL_geneID</u> : Homo sapiens gene ENSEMBL ID

<u>HUGO_dupGeneSymbol</u> : HUGO name for corresponding duplicated gene pair

<u>ENSEMBL_dupGeneID</u> : ENSEMBL ID for corresponding duplicated gene pair

<u>dup_branchLength</u> : duplication branch length calculated by TreeFam

<u>gene_familyName</u> : name of each family assigned by TreeFam

<u>gene_familySize</u> : number of genes in each family

**Table S2.** Evolutionary annotation of paralogous genes.

<u>WGD</u> : duplicated gene pair originated from Whole Genome Duplication events (WGD)

<u>SSD</u> : duplicated gene pair originated from Small Scale Duplication (SSD)

<u>ySSD</u> : duplicated gene pair originated from SSD event younger than WGD events

<u>wSSD</u> : duplicated gene pair originated from SSD event around WGD events

<u>oSSD</u> : duplicated gene pair originated from SSD event older than WGD events

**Table S3.** Differential gene expression analysis (DGE) between pairs of CNS regions.

This table describes the results of the differential gene expression analysis between all pairs of regions of the Central Nervous System.

<u>CNS_region_ref</u> : Central Nervous System region used as reference condition for DGE analysis

<u>nbDEG_dup</u> : number of differentially expressed genes (DEG) among duplicated genes

<u>nbDEG_singl</u> : number of DEG among singletons

<u>chi2_test</u> : Pvalue from chi-squared test

<u>DEG_dup_ratio</u> : number of DEG among duplicated genes divided by the total number of duplicated genes

<u>DEG_singl_ratio</u> : number of DEG among singletons divided by the total number of singletons

<u>chi2_Bonferroni</u> : P-value from chi-squared test corrected with Bonferroni

<u>odds_ratio</u> : occurrence of DEG among duplicates divided by the occurrence of DEG among singletons

<u>CNS_region_test</u> : Central Nervous System region used as tested condition for DGE analysis

**Table S4.** Tau score for region-specific genes.

This table lists the associations between genes and Central Nervous System regions according to their region-specific expression estimated using the Tau score.

<u>ENSEMBL_geneID</u> : Homo sapiens gene ENSEMBL ID

<u>Tau_score</u> : Tau score estimating region-specific expression of each gene

<u>CNS_region</u> : name of the Central Nervous System region

**Table S5.** Proportion of expressed genes in seven Central Nervous System (CNS) regions.

This table indicates the proportion of genes expressed for each of the seven regions of the Central Nervous System.

<u>CNS_region</u> : name of each Central Nervous System region

<u>Protein_coding_genes (%)</u> : percentage is with respect to the 16427 protein-coding genes expressed in CNS regions.

<u>Paralogous_genes (%)</u> : percentage is with respect to the 10335 paralogous genes expressed in CNS regions.

**Table S6.** Proportion of region-specific genes in seven Central Nervous System (CNS) regions.

This table indicates the proportion of region-specific genes for each of the seven regions of the Central Nervous System.

<u>CNS_region</u> : name of each Central Nervous System region

<u>Protein_coding_genes_with_region-specific_expression (%)</u> : percentage is with respect to the 2829 region-specific protein-coding genes expressed in CNS regions.

<u>Paralogous_genes_with_region-specific_expression (%)</u> : percentage is with respect to the 1985 region-specific paralogous genes expressed in CNS regions.

**Table S7.** WGCNA gene co-expression modules.

This table lists the gene composition of each co-expression module identified by Weighted Gene Correlation Network Analysis (WGCNA) methodology.

<u>WGCNA_moduleName</u> : label assigned by WGCNA to each gene co-expression module

<u>gene_list</u> : list of protein-coding ENSEMBL gene ID belonging to each module

<u>gene_nb</u> : number of protein-coding ENSEMBL genes belonging to each module

**Table S8.** WCGNA gene co-expression modules enriched for Gene ontology (GO) terms.

This table describes the results of the Gene Ontology (GO) terms over-representation analyses performed on each WGCNA co-expression module.

<u>WGCNA_moduleName</u> : label assigned by WGCNA to each gene co-expression module

<u>gene_nb</u> : number of protein-coding ENSEMBL genes belonging to each module

<u>GO_ID_MF</u> : Gene Ontology (GO) ID related to the most enriched Molecular Function (MF) term

<u>Pvalue_MF</u> : Pvalue assigned to the most enriched MF term. Pvalues lower than the Bonferroni correction value of 6.17E-04 are colored in red.

<u>oddsRatio_MF</u> : odds ratio assigned to the most enriched MF term

<u>GO_term_MF</u> : description of the most enriched MF term

<u>GO_ID_BP</u> : Gene Ontology (GO) ID related to the most enriched Biological Process (BP) term

<u>Pvalue_BP</u> : P-value assigned to the most enriched BP term. P-values lower than the Bonferroni correction value of 6.17E-04 are colored in red.

<u>oddsRatio_BP</u> : odds ratio assigned to the most enriched BP term

<u>GO_term_BP</u> : description of the most enriched BP term

**Table S9.** WGCNA gene co-expression modules corresponding to homogeneous gene families.

This table lists homogeneous gene families for which at least 60% of its member genes are included into the same WGCNA co-expression module.

<u>WGCNA_moduleName</u> : label assigned by WGCNA to each gene co-expression module

<u>gene Family_name</u> : name of each gene family found by TreeFam and considered as homogeneous family

**Table S10.** Pathway over-representation tests from homogeneous and heterogeneous gene families.

This table describes the results of the Reactome pathway over-representation analyses performed on homogeneous and heterogeneous gene families.

<u>reference_gene_list</u> : the number of genes from Homo sapiens PANTHER database that map to a particular pathway

<u>analyzed_gene_list</u> : the number of genes from homogeneous or heterogeneous families that map to a particular pathway

<u>expected_value</u> : the number of genes expected in the <u>analyzed_gene_list</u> for a particular pathway, based in the <u>reference_gene_list</u>

<u>fold_enrichment</u> : the fold enrichment of the genes observed in the <u>analyzed_gene_list</u> over the <u>expected_value</u>

<u>raw_Pvalue</u> : the raw P-value calculated by Fisher exact test. It is the probability that the number of genes observed in the <u>analyzed_gene_list</u> for particular pathway occurred by chance, as determined by the <u>reference_gene_list</u>

<u>FDR</u> : the False Discovery Rate calculated by the Benjamini-Hochberg procedure. A threshold of 0.05 is used to filter results, so all over-represented pathways shown are valid for an overall FDR<0.05.

**Table S11.** Patient GTEx ID to CNS region name.

This table indicates the correspondence between the patient's GTEx ID and the associated CNS regions.

<u>Patient_GTEX_ID</u> : the list of GTEx patient sample ID used in our study

<u>CNS_regions_name</u> : the CNS region associated to each patient sample

**Table S12.** Singleton gene list.

This table lists the genes annotated as expressed singletons in our study.

<u>Singleton_genes</u> : the human ENSEMBL ID associated to each gene

**Table S13.** Homogeneous gene list.

This table lists the genes belonging to homogeneous families in our study.

<u>Genes_from_homogeneous_families</u> : the human ENSEMBL ID associated to each gene

**Table S14.** Heterogeneous gene list.

This table lists the genes belonging to heterogeneous families in our study.

<u>Genes_from_heterogeneous_families</u> : the human ENSEMBL ID associated to each gene

**Table S15.** WGCNA modules and region-specific genes.

This table indicates for each WGCNA co-expression module the proportion of region-specific genes and their associated number of regions.

<u>WGCNA_moduleName</u> : label assigned by WGCNA to each gene co-expression module

<u>gene_nb</u> : number of protein-coding ENSEMBL genes belonging to each module

<u>perc_regionSpecificGenes</u> : proportion of region-specific genes

<u>nb_associatedRegions</u> : number of regions associated with region-specific genes

**Table S16.** Multivariate linear regression to model the relationship between the region-specificity of genes (corresponding to the Tau score of genes) and explanatory variables (expression, duplication status, the age of the duplication and the type of the duplication) depending on the reference group. Linear models were done in parallel on the set of protein coding genes in all of the paper selected by a threshold of maximal expression > 0.1 RPKM and with a restricted set of genes selected by a threshold of maximal expression > 1 RPKM.

**Table S17.** Enrichments in genes from region-specific families (in the same region) for the tested and reference gene groups.

**Table S18.** Enrichments in brain disorder-associated genes for the tested and reference gene groups.

**Table S19.** List of brain disorder-associated genes.

<u>ENSEMBL_geneID</u> : Homo sapiens gene ENSEMBL ID

<u>HUGO_geneSymbol</u> : Homo sapiens gene name based on HUGO nomenclature

**Table S20.** Enrichments in gene pairs according to genomic distance.

**Table S21.** Enrichments in region-specific genes for the tested and reference gene groups (selected genes with a threshold of maximal expression > 1 RPKM).

